# Bimanual Postural Similarity Facilitates Haptic Memory-Guided Reaching

**DOI:** 10.1101/2025.10.27.684813

**Authors:** Ivan Camponogara, Robert Volcic

**Author notes:** **Corresponding author information:** Ivan Camponogara, Department of Psychology, College of Natural and Health Sciences, Zayed University, Abu Dhabi, United Arab Emirates.

## Abstract

In the visual domain, the precision of memory-guided reaching movements gradually decays as the delay between vision withdrawal and movement onset increases. Such a decrease is often linked to the impact of visual memory decay and the sensory transformation of visual target positional information into haptic-specific coordinates. It is, however, unclear whether this holds for haptically encoded spatial locations, where the target and reaching arm positions are both defined through haptics. Here, we asked participants to perform memory-guided right-handed reaching movements toward targets held with the left hand. In the Delay block, the left hand was removed after 2 seconds, and a right-handed reach was performed after a variable delay of 2, 4, or 6 seconds. In the No-delay block, the left hand was on the target for the entire duration of the trial. Targets were positioned in front (Center) and to the Left of the participants, such that the posture of the left arm during encoding was either the same or different from the final posture of the contralateral reaching arm. The introduction of the shortest delay caused a decrease in accuracy and precision, indicating that on-line haptic inputs play a crucial role in guiding reaching movements. However, performance did not worsen with longer delays, particularly when the two arms were in similar postures. Our findings suggest that postural information may contribute to guiding actions when on-line haptic feedback is unavailable.

**Highlights:**

- Memory-guided reaching accuracy in the visual domain declines as the delay between vision and action increases.
- This study tested whether similar effects occur for haptically encoded spatial locations using right-hand reaches toward left-hand-held targets.
- Introducing short delays (2 s) reduced accuracy and precision, highlighting the importance of on-line haptic input for movement guidance.
- Longer delays (4–6 s) did not further impair performance, particularly when arm postures during encoding and reaching were similar.
- Postural information supports reaching accuracy when real-time haptic feedback is unavailable.

## 1 Introduction

In visually guided reaching, the continuous real-time availability of target visual information is fundamental for the execution of precise and accurate movements (Adamovich, Berkinblit, Smetanin, Fookson, & Poizner, 1994; Berkinblit, Fookson, Smetanin, Adamovich, & Poizner, 1995; Elliott, 1988; Heath & Binsted, 2007; Heath, Westwood, & Binsted, 2004; M. Khan et al., 2006; M. A. Khan & Franks, 2000; Krigolson & Heath, 2004; Lacquaniti et al., 1997; Lemay, Gagnon, & Proteau, 2004; Lemay & Proteau, 2001, 2002; McIntyre, Stratta, & Lacquaniti, 1998; Rolheiser, Binsted, & Brownell, 2006; Smyrnis, Gourtzelidis, & Evdokimidis, 2000; Soechting & Flanders, 1989a, 1989b; Westwood, Heath, & Roy, 2001, 2003). Blocking vision at least two seconds prior to initiating a movement negatively impacts action performance, leading to a consistent reduction in both precision and accuracy (Elliott, 1988; Heath & Binsted, 2007; Heath et al., 2004; Lacquaniti et al., 1997; Monaco et al., 2010; Pettypiece, Culham, & Goodale, 2009; Smyrnis et al., 2000; Westwood et al., 2001, 2003). The underlying reason for this decline may be related to the effect of a visual memory deterioration on the transformation of target visual inputs into haptic information. When reaching for a visually sensed target, a cross-sensory transformation needs to occur: the visual representation of the target is transformed into a haptically coordinated pattern of arm movements to guide the hand (Darling & Miller, 1993; Flanders, Tillery, & Soechting, 1992; Prablanc, Pelisson, & Goodale, 1986; Soechting & Flanders, 1989a, 1989b; Tillery, Flanders, & Soechting, 1991). This process is affected by intrinsic noise, as sensory inputs from one modality need to be cross-transformed in another sensory modality (Tagliabue & McIntyre, 2014). Since visual memory rapidly decays over time (Heath & Binsted, 2007; Heath et al., 2004; M. Khan et al., 2006; Krigolson & Heath, 2004; Rolheiser et al., 2006; Westwood et al., 2001, 2003), the combination of the memory decay with the cross-sensory transformation process leads to inaccurate and imprecise actions (Berkinblit et al., 1995; Darling & Miller, 1993; Soechting & Flanders, 1989a, 1989b; Tillery et al., 1991).

There is, however, a debate about whether the same holds for haptically defined target positions, as it is still unclear whether the haptic memory of a previously sensed target can be used to perform a successful reach. While visual target information needs to be cross-transformed into haptic information, coding of the target position in the haptic domain can occur through the direct encoding of arm postures (Proske & Gandevia, 2012). These can be used to directly match the contralateral arm posture to successfully reach the target (Wrisberg & Winter, 1985). Thus, in memory-guided reaching, directly encoding target positional information through arm posture could lead to a more robust storage of this information which could persist for a relatively longer period. The encoding and storage of haptic inputs could lead to a similar action performance across delays. However, studies exploring action guidance under haptic memory showed contrasting results.

Studies comparing actions performed either with or without a fixed time delay consistently and, unsurprisingly, reported a worse action performance in memory-guided reaching (Berkinblit et al., 1995; Darling & Miller, 1993; Jones, Fiehler, & Henriques, 2012; Jones & Henriques, 2010; Pettypiece et al., 2009; Soechting & Flanders, 1989a, 1989b; Tillery et al., 1991). A decrease in action performance is indeed expected whenever on-line haptic inputs are prevented. Critically, the above mentioned studies lack a comparison between actions performed at different time delays; a crucial contrast to define whether the encoding and comparison of arm positions through haptics could dampen the decrease in action performance usually seen in memory-guided reaching. In this regard, studies comparing action performance across delays showed incongruent results. Some studies showed that a delay of 5 and 45 seconds (Wrisberg & Winter, 1985) or 1, 2, or 10 seconds (Chapman, Heath, Westwood, & Roy, 2001) between the sensing and the reaching phases does not affect action performance: actions were equally accurate and precise regardless of the delay. In contrast, Stelmach (1969) and Stelmach and Walsh (1972) showed a lower accuracy and precision when the delay between sensing and reaching phases was 15 compared to 50 seconds (Stelmach, 1969) or 5 compared to 20 seconds (Stelmach & Walsh, 1972), with a plateau in precision at delays longer than 15 seconds (Stelmach, 1969). Thus, it is still unclear whether the use of haptic postural information from the arm holding the target could prevent a decay in action performance in memory-guided reaching.

Here, we shed light on how memory-guided reaching unfolds in the haptic domain by systematically comparing actions performed without delay to those executed after varying temporal delays following a haptic sensing phase. In a No-delay block, participants were asked to hold a target for two seconds and reach for it with the contralateral hand. In a Delay block, participants were asked to retract the sensing hand 2, 4, or 6 seconds before reaching for the target with the other hand. If haptic memory-guided reaching consists of replicating the posture of the arm holding the target by the reaching hand, we expect a small impact of the delay on the action accuracy and precision within the Delay block. In contrast, if haptic postural information from the hand holding the target is further transformed before being used by the reaching arm, we expect a dramatic decrease in action accuracy and precision within the Delay block.

In an additional experimental manipulation, we explored whether the postures of the arm holding the target and of the reaching arm are directly encoded through a direct comparison between joint configurations. The manipulations consisted of modulating the positions of the targets, which were located either at the Center or to the Left (45 degrees) of the participants’ torso. For each direction, we used two distances, one Near (150 mm) and one Far (300 mm). When reaching a handheld target aligned with the center of the body, the final posture of the reaching arm mirrors that of the arm holding the target. In contrast, when reaching a handheld target positioned laterally, the sensing and reaching arms have two different postures. If action performance relies on a direct comparison between arm postures, we expect a decrease in performance for targets placed on the Left compared to those located at the Center. If not, a similar action performance across target positions (Left and Center) is expected.

## 2 Experimental Procedures

### 2.1 Participants

We tested thirty participants (6 left-handed, 14 males, age 19.6*±*1.27 years). All had normal or corrected-to-normal vision and no known history of neurological disorders. All participants were naïve to the purpose of the experiment and were provided with a subsistence allowance for their participation. The experiment was undertaken with the understanding and informed written consent of each participant, and the experimental procedures were approved by the Institutional Review Board of New York University Abu Dhabi.

### 2.2 Apparatus

The set of stimuli consisted of four 15 mm high 3D-printed cylinders with a diameter of 80 mm positioned at 150 mm (i.e., Near) and 300 mm (i.e., Far) from a start position. Two targets were aligned with the center of the participant’s torso, whereas two were at 45 degrees to the left of the participant (Figure 1 A). The start position was a 3D printed cylinder (35 mm height, 20 mm diameter), so-called “base”, which supported a second cylinder called “marker” (40 mm height, 15 mm diameter). A pure tone of 1000 Hz, 100 ms duration was used to signal the start of the trial, while a tone of 500 Hz of the same duration was used to signal its end. The centroid of the marker object was acquired on-line at 200 Hz with sub-millimeter resolution using an OptiTrack system (NaturalPoint, Inc.) by defining a rigid body using three reflective markers placed on the top of the cylinder.

**Figure 1.**
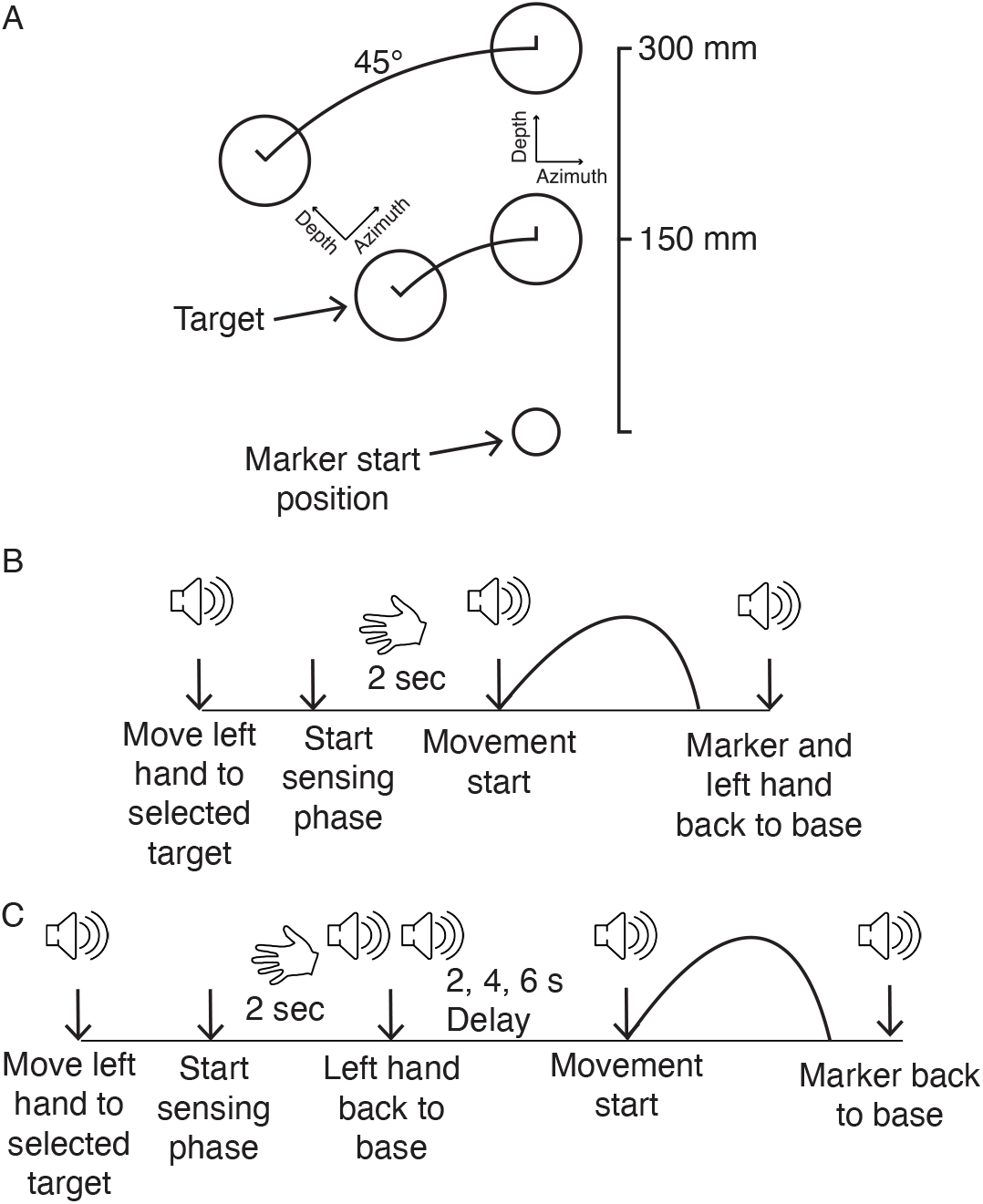
A) Experimental setup. Targets were positioned at 150 mm (i.e., Near) and 300 mm (i.e., Far) from the start position, where the marker cylinder was positioned. Two targets were aligned with the center of the participant’s torso, whereas two were at 45 degrees to the left of the participant, but always at 150 mm (i.e., Near) and 300 mm (i.e., Far) from the start position. B-C) Representation of the task in the No-delay and Delay blocks. In both blocks, the participants were instructed to move the left hand to one of the four targets at the beginning of each trial. B) In the No-delay block after a two-second sensing phase, a start tone was delivered, and participants moved the marker to the held target. C) In the Delay block, the two-second sensing phase was followed by two consecutive sounds, which signaled participants to move the left hand back to the base. After a variable delay from the consecutive sounds (2, 4, or 6 seconds) the start tone was delivered, and participants moved the marker to the remembered target position. In both blocks, a tone signaled the participants to move the marker back to the base after movement completion.

### 2.3 Procedure

Participants were blindfolded for the whole duration of the experiment. Two blocks were performed: a No-Delay block and a Delay block. In each block, each trial began with participants holding the marker cylinder between the index and thumb of the right hand, and the base cylinder between the index and thumb of the left hand. A pre-recorded voice then signaled the participant to move the left hand to a randomly selected target (i.e., Center Far, Center Near, Left Far, Left Near). The contact of the left hand with the target started a “sensing” phase, where participants sensed the target position for two seconds. Subsequently, in the No-delay block, at the start tone, participants were required to move the marker to the center of the held target as accurately and precisely, and as fast as possible (Figure 1B). In the Delay block, instead, the two-second sensing phase was interrupted by two consecutive sounds (1000 Hz, 100 ms duration, 200 ms separation between beeps), which signaled the participants to move the left hand back to the base. Then, the start tone was randomly delivered after 2, 4, or 6 seconds and participants were instructed to move the marker to the remembered target location as accurately and precisely, and as fast as possible (Figure 1C). In both blocks, the end tone was delivered two seconds after the start tone, and participants moved the marker (and the left hand in the No-delay block) back to the base. Before each block, participants underwent a practice phase consisting of 15 trials per target, where they got accustomed to the task. We run 15 trials for each target and delay, leading to 60 trials for the No-delay and 180 trials for the Delay block. Three five-minute breaks were introduced every 60 trials in the Delay block. For each participant, the Delay block was run first to avoid any learning effect.

### 2.4 Data analysis

Kinematic data were analyzed in R (R Core Team, 2024). The raw data were smoothed and differentiated with a third-order Savitzky-Golay filter with a window size of 21 points. These filtered data were then used to compute velocities and accelerations of the marker in three-dimensional space. Movement onset was defined as the moment of the lowest, non-repeating marker acceleration value prior to the continuously increasing marker acceleration values, while the end of the movement was defined by applying the same algorithm, but starting from the end of the recorded values (Camponogara & Volcic, 2021a). From the 7200 trials we discarded from further analysis the trials in which the end of the movement was not captured correctly or in which the missing marker samples could not be reconstructed using interpolation. The exclusion of these trials (483 trials, 6.7% of all trials) left us with 6717 trials for the final analysis. For each trial, we calculated the endpoint positions along the depth and azimuth axes according to the movement direction (Figure 2)

**Figure 2.**
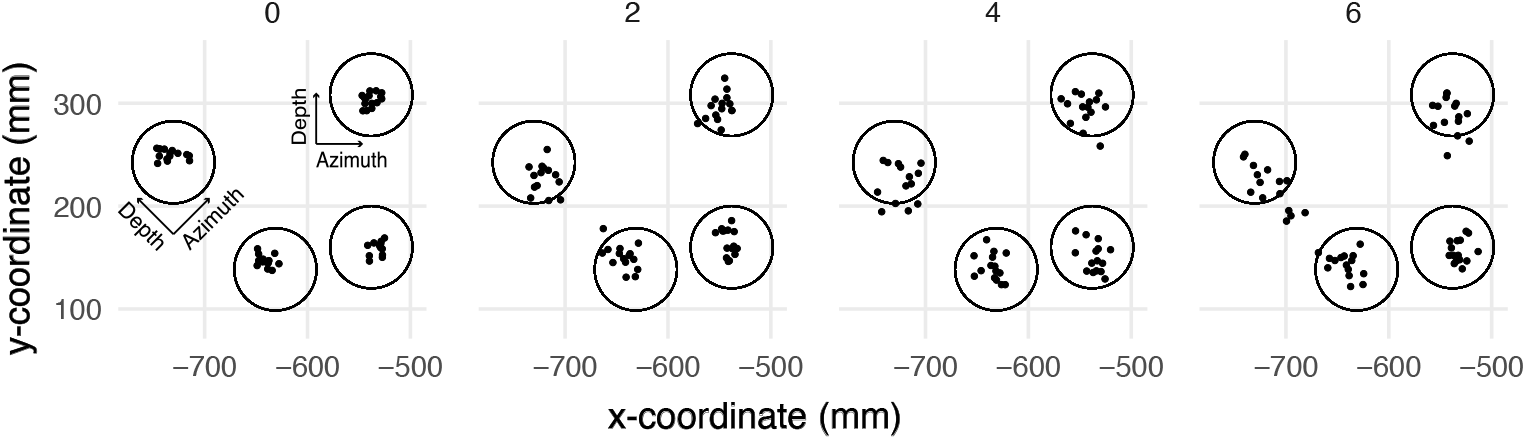
Final landing points of a representative participant in the No-delay (0 seconds) and Delay blocks (2, 4, 6 seconds). Each panel shows the landing positions of the 60 trials in each block (15 trials for each target).

We focused our analyses on the endpoint errors, defined as the distances between the center of the target and the marker’s landing point along the depth and azimuth axes. We analyzed these data using a Bayesian multivariate mixed-effects distributional regression model, estimated using the *brms* package (Bürkner, 2017), which implements Bayesian multilevel models in R using the probabilistic programming language Stan (Carpenter et al., 2017). Unlike conventional regression models that assume a constant standard deviation across observations, a distributional regression model estimates not only the location parameters but also the standard deviations of the location parameters. The Movement Direction (Center, Left), Delay (0, 2, 4, 6), and Target Position (Near and Far) were encoded in a unique categorical variable Condition, which was used as the fixed-effect of our model (i.e., Center 0 Near, Center 2 Near, etc.). Thus, the estimates of the Condition parameters (*β*_*Condition*_) correspond to the average endpoint error (i.e., accuracy) along the azimuth and depth axes of each Movement Direction, Delay and Target Position, whereas the *σ*_*Condition*_ parameters correspond to the within-participant endpoint variability (i.e., precision) in azimuth and depth. The model included the independent random (group-level) effects for subjects. The model was fitted considering weakly informative prior distributions for each parameter to provide information about their plausible scale. We used Gaussian priors for the Condition fixed-effect predictor (*β*_*Condition*_: mean = 0 and sd = 50, *σ*_*Condition*_: mean = 0 and sd = 5), whereas for the group-level standard deviation parameters we used zero-centered Student *t* -distribution priors (azimuth = *df* = 3, scale = 15, depth = *df* = 3, scale = 19). Finally, we set a prior over the correlation and residual correlation matrix that assumes that smaller correlations are slightly more likely than larger ones (LKJ prior set to 2).

We ran four Markov chains simultaneously, each for 4,000 iterations (1,000 warm-up samples to tune the MCMC sampler) with the delta parameter set to 0.99 for a total of 12,000 post-warm-up samples. Chain convergence was assessed using the 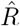statistic (all values equal to 1) and visual inspection of the chain traces. Additionally, predictive precision of the fitted models was estimated with leave-one-out cross-validation by using the Pareto Smoothed importance Sampling (PSIS). All Pareto k values were below 0.5.

The obtained posterior distributions represent the probabilities of the parameters conditional on the priors, model, and data, and they represent our belief that the “true” parameter lies within some interval with a given probability. We summarize these posterior distributions by computing the medians and the 95% Highest Density Intervals (HDI). The 95% HDI specifies the interval that includes, with a 95% probability, the true value of a specific parameter. To evaluate the differences in accuracy and precision between two conditions, we simply subtracted the posterior distributions of *β*_*Condition*_ and *σ*_*Condition*_ between specific conditions. The resulting distributions are denoted as the credible difference distributions and are again summarized by computing the medians and the 95% HDIs.

For statistical inferences about the *β*_*Condition*_ and *σ*_*Condition*_ we assessed the overlap of the 95% HDI with zero. A 95% HDI that does not span zero indicates that the endpoint error (*β*_*Condition*_) or endpoint variability (*σ*_*Condition*_) in depth and azimuth was credibly different than zero in that specific condition. For statistical inferences about the differences between conditions, we applied an analogous approach. A 95% HDI of the credible difference distribution that does not span zero is taken as evidence that the endpoint error (*β*_*Condition*_) or endpoint variability (*σ*_*Condition*_) in the two conditions differ from each other.

From here onward, we will use the term “accuracy” when reporting the results of *β*_*Condition*_ and “precision” when reporting the results of *σ*_*Condition*_.

## 3 Results

To structure the presentation of our findings, we first consider accuracy across the different conditions of the task, and subsequently turn to precision.

### 3.1 Accuracy

In general, participants overshot the Near targets (Figure 3A) and they undershot the Far targets (Figure 3C). Moreover, the accuracy in azimuth was modulated according to movement direction: the marker landed on the right side of the targets positioned on the Left and on the left side of the targets positioned in the Center (Figure 3B and D).

**Figure 3.**
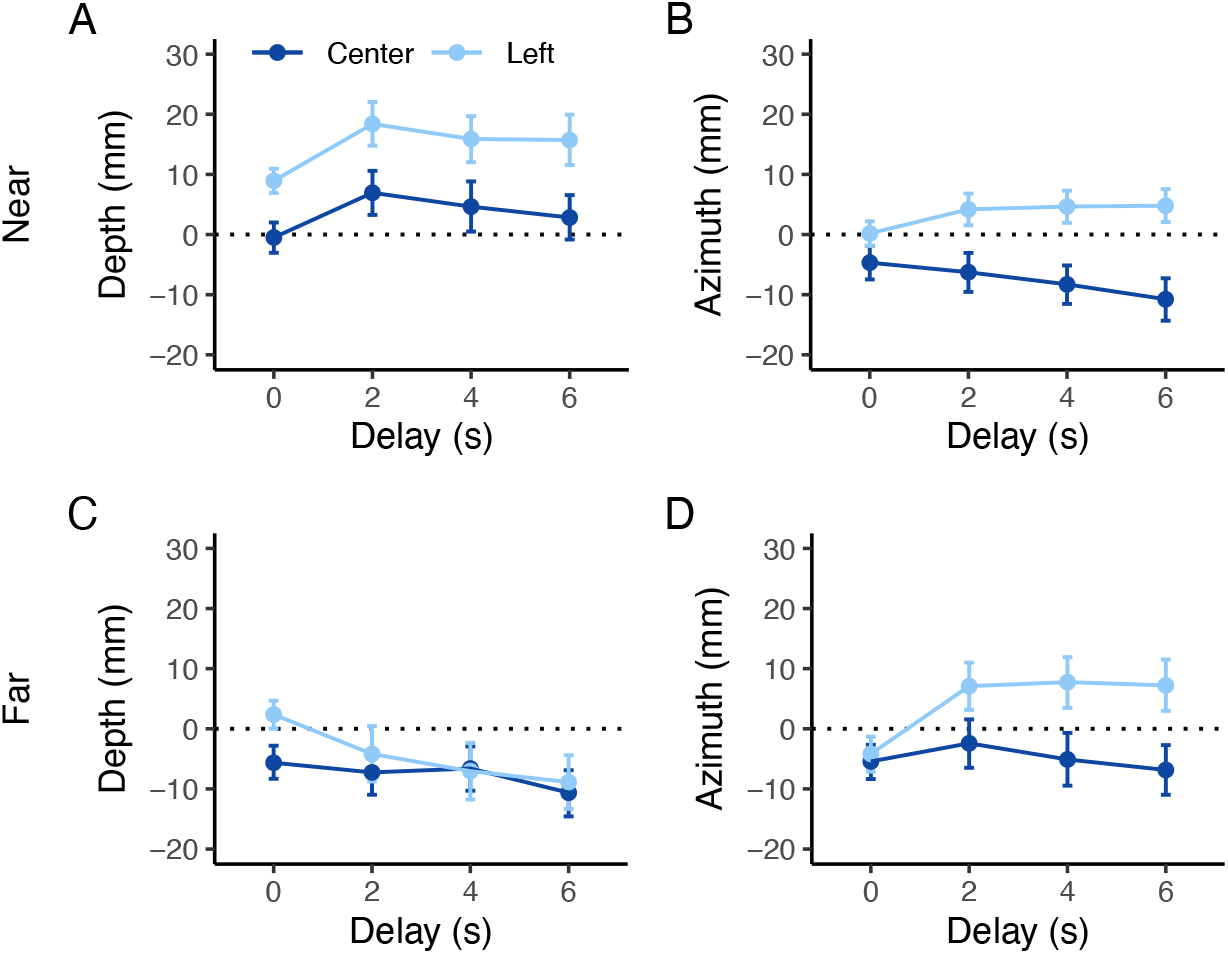
Estimates of the accuracy in depth and azimuth of all the conditions and delays for the Near (A, B) and Far (C, D) targets. The points represent the median and the error bars denote the 95% HDI. The dashed line represents the center of the targets. Positive values in depth indicate an overshoot, whereas positive values in azimuth indicate that the marker landed on the right side of the target.

#### 3.1.1 Effect of Delay

Center movements were generally longer in the delay block compared to the No-delay block when the target was Near (Figure 4A, 0-2 comparison), but not when it was Far. We found a similar pattern for Left movements, with the exception of a slight undershoot for the Far target when a delay was introduced (Figure 4C, 0-2 comparison). In contrast, while accuracy in the azimuth direction was similar between the delay and no delay blocks for the Center movements, it showed a modulation for the Left movements according to the target distance: it was lower in the Delay than in the No-delay block when the target was Far (Figure 4D, 0-2 comparison), but not when it was Near (Figure 4B, 0-2 comparison). Interestingly, increasing the delay from 2 s to 4 s or even to 6 s did not produce further changes in accuracy for both the Center and Left movements (Figure 4, 2-4 and 4-6 comparisons). These results suggest that haptic inputs for action performance can be used up to 6 seconds following the sensing phase. However, haptic memories of targets positioned at the Center are stored longer than those positioned laterally, as generally (except for the movements toward Near targets in the depth direction), no differences were found between the delay and No-delay blocks for the Center targets.

**Figure 4.**
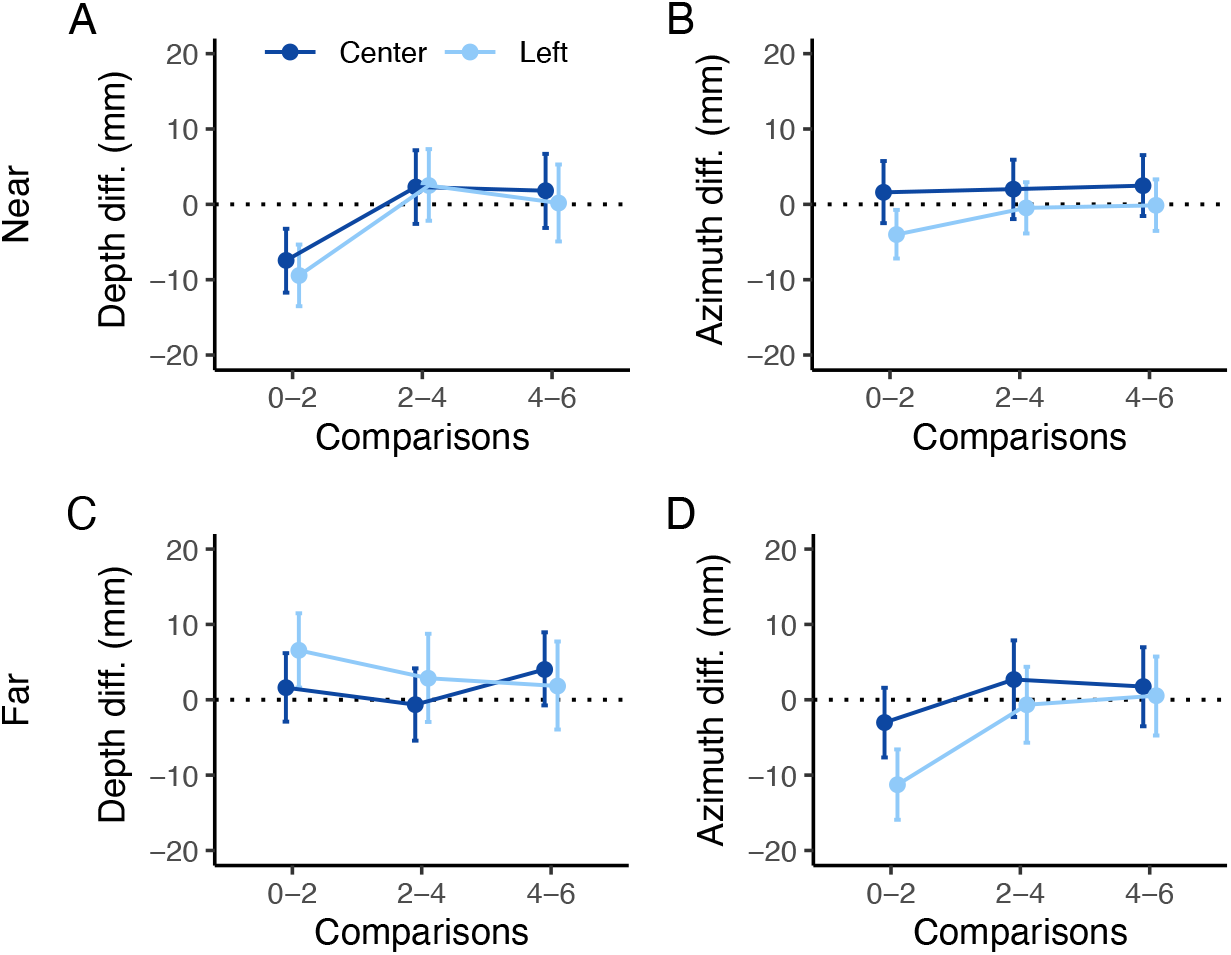
Estimates of the differences in accuracy in depth and azimuth (left and right columns, respectively) for the Near and Far targets (upper and lower rows, respectively). The points represent the median and the error bars denote the 95% HDI of the difference between estimates. The dashed line represents the points of equal accuracy between pairs of delays. A, B) Difference in accuracy between delays for the Near target within each movement direction (Center and Left); C, D) Difference in accuracy between delays for the Far target within each movement direction (Center and Left). Positive values in depth difference indicate a longer reaching in the first delay compared to the second, whereas positive values in azimuth difference indicate a more rightward reaching in the first delay compared to the second. Comparisons are indicated on the x-axis.

#### 3.1.2 Effect of Movement Direction

We found a general bias in accuracy according to movement direction. Movements toward the Center Near targets ended generally closer (in depth) than those toward the Left Near targets in both the No-delay and Delay blocks. For Far targets, this holds only for the No-delay block, whereas for the Delay block, a similar accuracy was found between Center and Left directions (Figure 5A). Additionally, movements in the Center direction were directed more leftward compared to those to the Left, except for Far targets in the No-delay block (Figure 5B).

**Figure 5.**
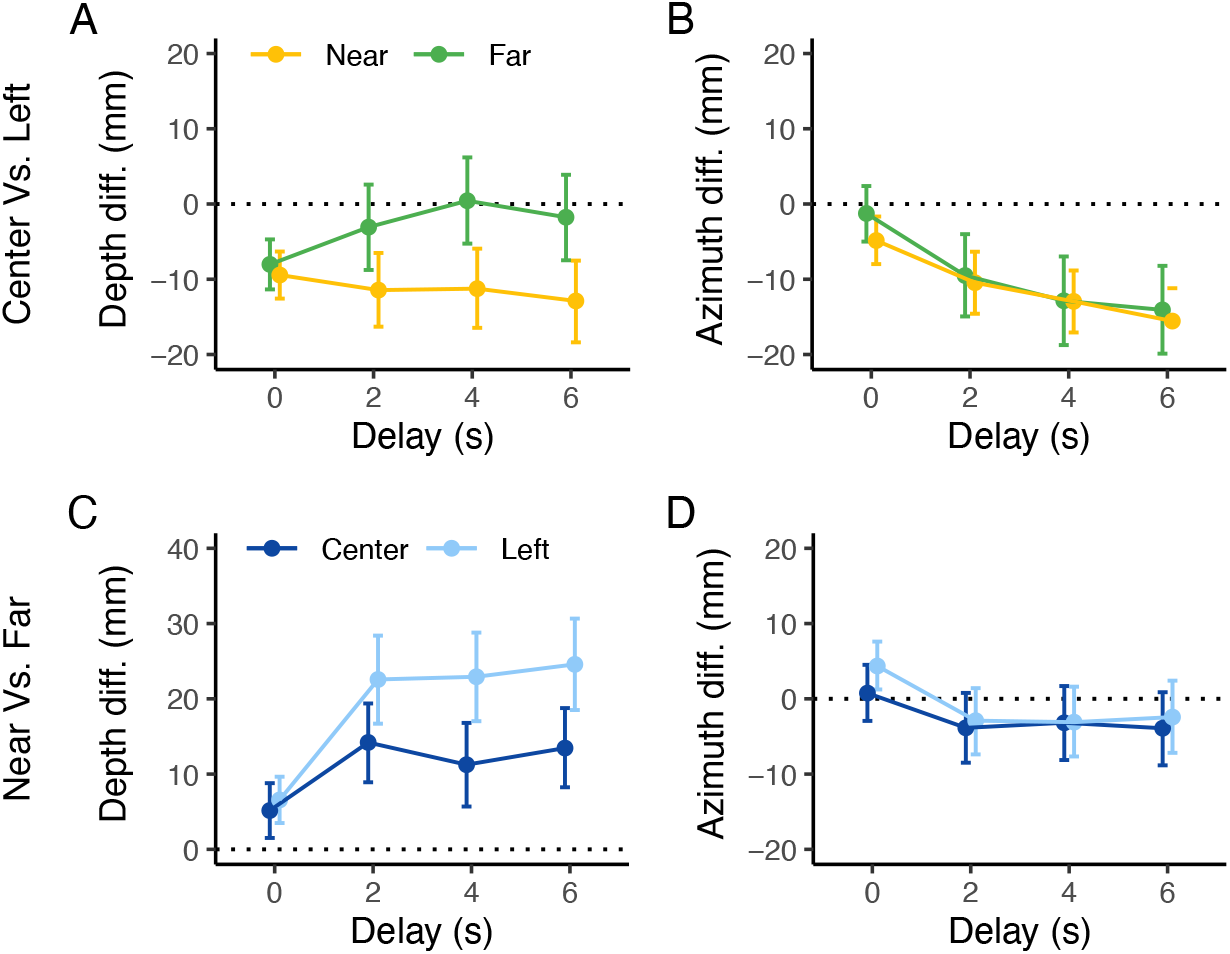
Estimates of accuracy differences in depth and azimuth. The points represent the median, and the error bars denote the 95% HDI. A, B) Differences between Center and Left movement directions for each target distance and for each delay. Negative values in depth indicate shorter movements when reaching the Center compared to the Left targets. In contrast, negative values in azimuth indicate that Center targets were reached more leftward than Left targets. C, D) Differences in accuracy between Far and Near targets for each movement direction and for each delay. Positive values in depth difference indicate a longer reaching for the Far compared to the Near targets, whereas positive values in azimuth difference indicate that Far targets were reached more rightward than Near targets.

#### 3.1.3 Effect of Target Distance

Generally, there was a tendency for a larger overshoot for the Near compared to the Far targets in both Delay and No-delay blocks and both movement directions (Figure 5C). Accuracy in azimuth was similar across delays and movement directions between Near and Far targets, with the exception of the Left movements in the No-delay block, where there was a tendency to land more rightward for the Near compared to the Far targets (Figure 5D).

### 3.2 Precision

In general, precision was higher (i.e., endpoint variability was lower) when actions were performed without delay compared to when a delay was introduced (Figure 6). Interestingly, precision was approximately constant for all three delay durations.

**Figure 6.**
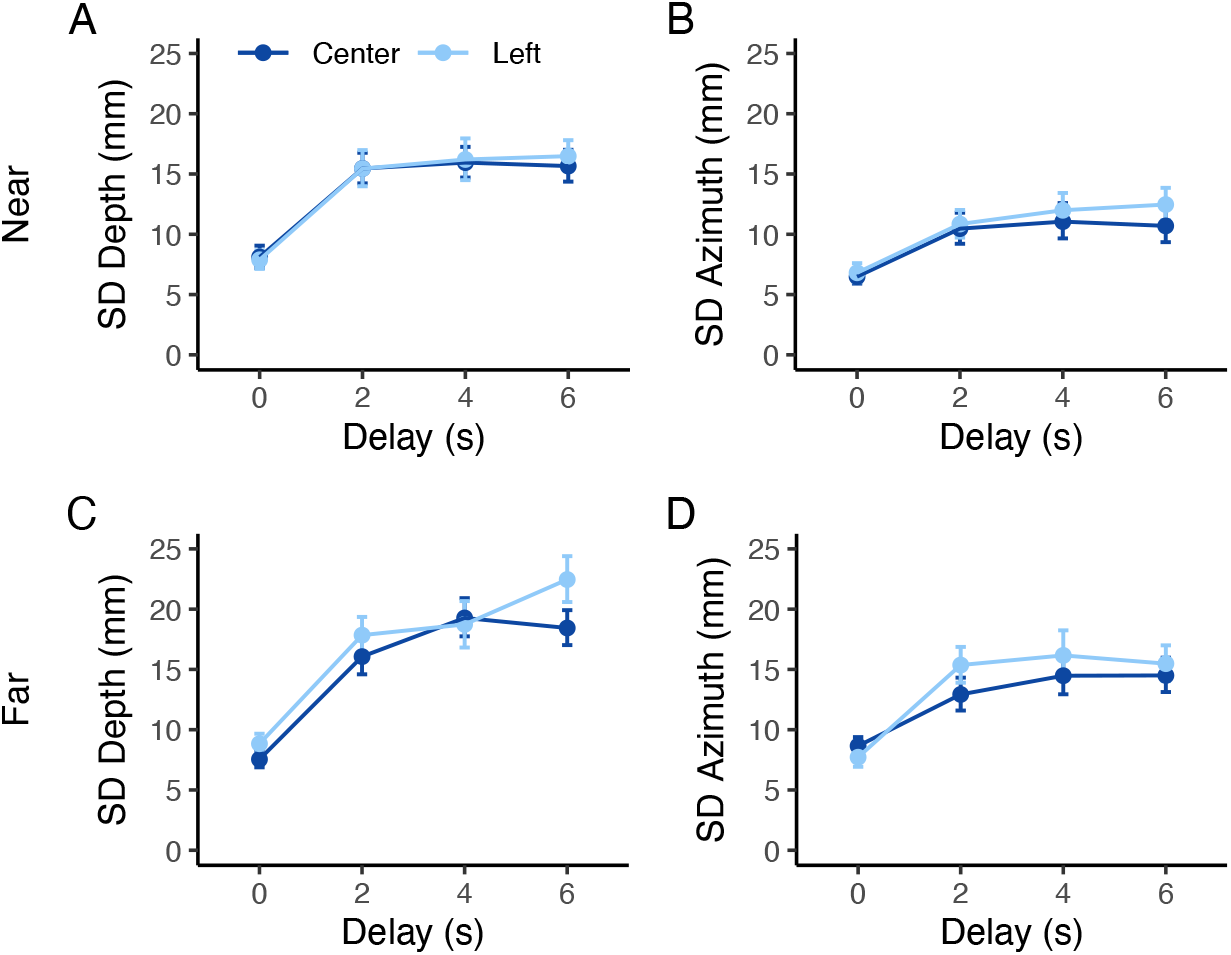
Estimates of the precision in depth and azimuth for all the conditions and delays for the Near (A, B) and Far (C, D) targets. The points represent the mean and the error bars denote the 95% credible intervals of the estimates.

#### 3.2.1 Effect of Delay

Introducing a delay between the sensing phase and the onset of movement generally reduced precision (Figure 7). However, precision was similar regardless of the delay for each target distance, with the exception of the 2-4 delay comparison for Center movements, and 4-6 comparison for the Left movements in depth (Figure 7C).

**Figure 7.**
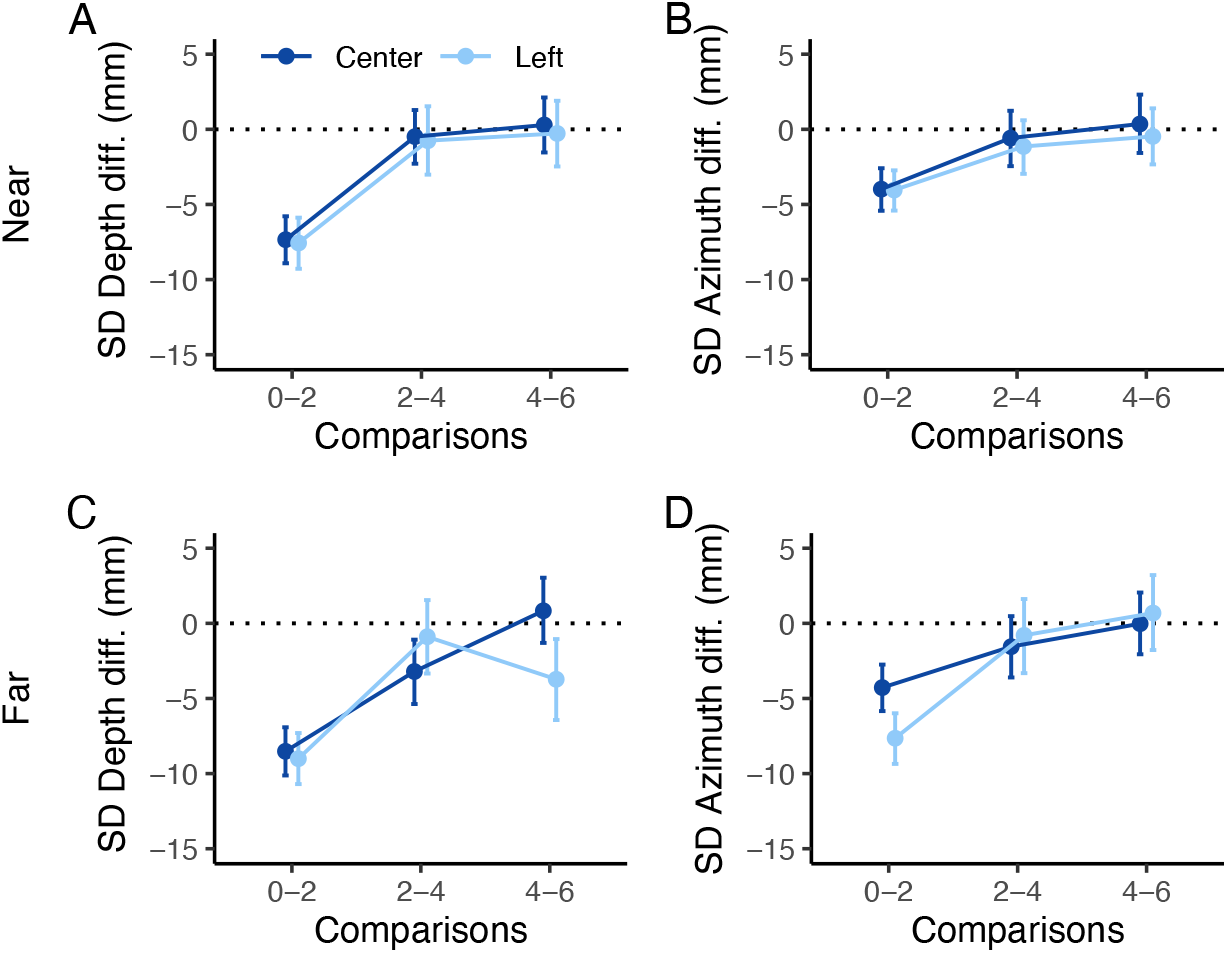
Estimates of the precision in depth and azimuth for all conditions and delays. The points represent the mean and the error bars denote the 95% credible intervals of the estimates. A, B) Difference in precision between delays for the Near target within each movement direction; C, D) Difference in precision between delays for the Far target within each movement direction. Positive values indicate a higher variability in the first delay compared to the second (comparisons are indicated on the x-axis).

#### 3.2.2 Effect of Movement Direction

Precision was generally similar for Center and Left directions regardless of the delay or target distance (Figure 8A and B). However, at 6 seconds delay, Center movements toward the Far target were less variable than Left movements along the depth axis (Figure 8A, 6 seconds delay). The same holds for the 2 second delay in azimuth (Figure 8B, 2 seconds delay).

**Figure 8.**
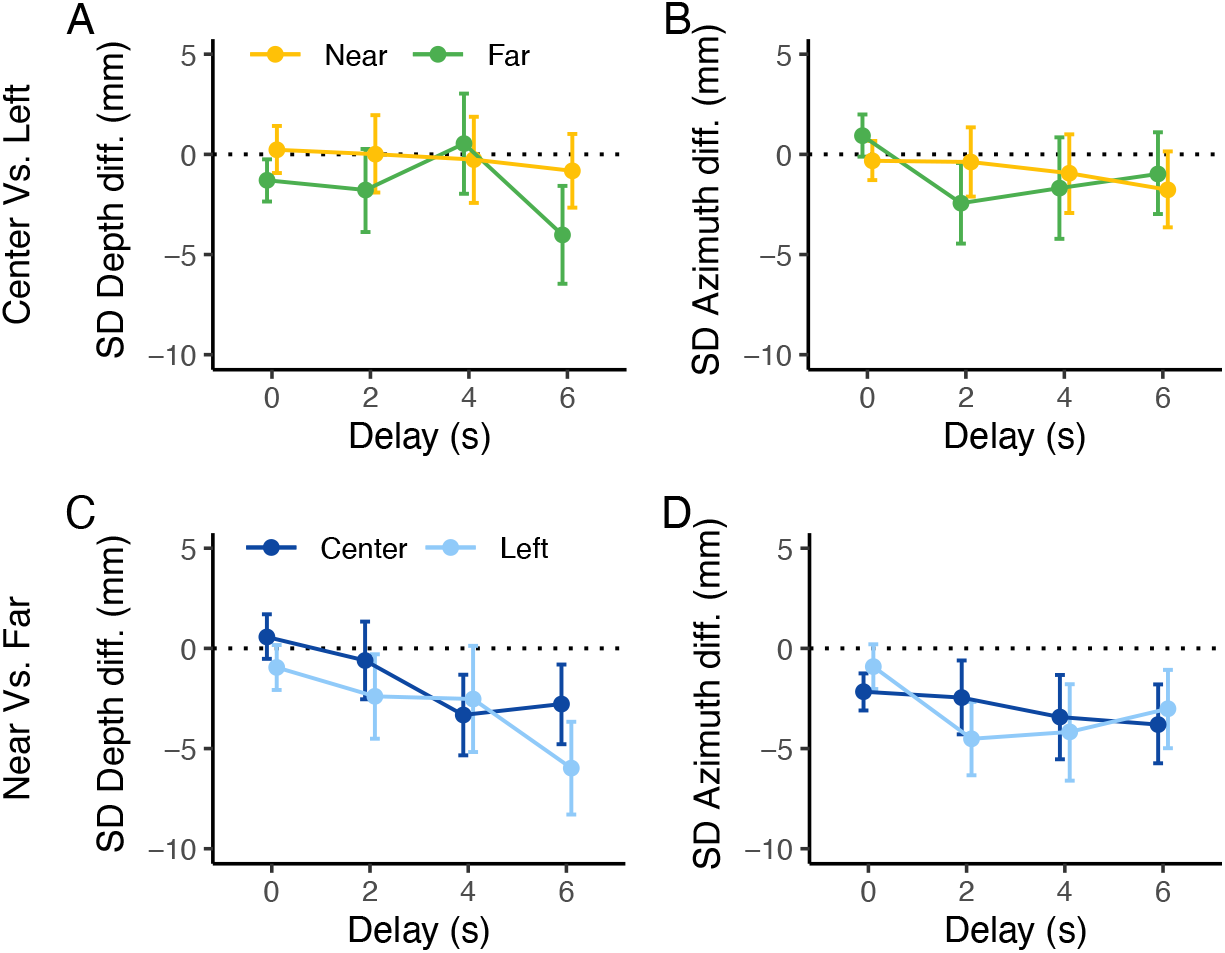
Estimates of the differences in precision in depth and azimuth (left and right columns, respectively). A, B) Differences between Center and Left movement directions for each target in each delay. Negative values indicate a lower variability in depth or azimuth in the Center compared to the Left movement direction for each target (Near and Far). C, D) Differences between Near and Far targets for each movement direction in each delay. Negative values indicate a lower variability in either depth or azimuth when reaching the Near target compared to the Far target in each movement direction (Center and Left).

#### 3.2.3 Effect of Target Distance

Generally, movements toward Near targets were less variable (i.e., more precise) than those toward Far targets in both depth and azimuth for Left movement direction in the Delay block (Figure 8C and D, light blue). For movements directed to the Center, the variability was lower for Near compared to Far targets at 4 and 6 seconds delay (Figure 8C, dark blue), whereas it was lower for Near compared to Far targets in all the blocks (Delay and No-delay) in azimuth (Figure 8D, dark blue).

## 4 Discussion

Here, we investigated the effect of haptic memory in reaching performance. Blindfolded participants were asked to use their left hand to sense a target positioned at the Center or at the Left (45 degrees) of their torso, either Near or Far from a starting position. Following a two seconds sensing phase, in a Delay block, participants were asked to retract the left hand and after 2, 4 or 6 seconds, reach the target with the contralateral (right) hand. In the No-delay block, instead, reaching was initiated immediately after the sensing phase, with the target being constantly held in the participants’ hand during reaching (Figure 1).

We found that, in general, introducing a delay modulates action accuracy, but primarily for targets on the left side of the body. The target location was more accurately stored for a longer time when the postures of the reaching and the sensing arms matched (i.e., when the targets were at the Center) than when they did not (i.e., when the targets were on the left). Our results also showed a decreased precision with the introduction of a delay between the sensing and reaching phases. However, precision was generally similar across delays and movement directions. This confirms previous results in the haptic domain that showed similar action performance across haptic delays (Chapman et al., 2001; Goble & Brown, 2008; Jones & Henriques, 2010). Taken together, our results suggest that replicating the posture of the sensing arm with the reaching arm could facilitate memory-guided actions toward a previously sensed haptic target, and that such postural information might guide actions when haptic on-line information is unavailable.

In visually guided reaching, action performance decreases as the delay between target sensing and movement onset increases (Heath & Binsted, 2007; Heath et al., 2004; Westwood et al., 2001, 2003). This is because visual information about the target needs to undergo a sensory transformation into haptic (proprioceptive and tactile) information to guide the reaching hand (McIntyre et al., 1998; Soechting & Flanders, 1989a, 1989b). Introducing a delay between the sensing and the movement onset phase dramatically affects the visual target positional information, disrupting the above-mentioned process (Berkinblit et al., 1995; Darling & Miller, 1993; Heath & Binsted, 2007; Heath et al., 2004; Soechting & Flanders, 1989a, 1989b; Tillery et al., 1991; Westwood et al., 2001, 2003). In contrast, in the haptic domain, target positional information may have been stored directly in arm postural information, without the involvement of any transformation process between modalities. As a result, the impact of increasing delays on accuracy and precision was minimal, especially when the arm postures were similar.

The role of haptics is to provide continuous information about the magnitude of limb/joints extensions (Proske & Gandevia, 2012), which, in bimanual object manipulation (i.e., passing an object from one hand to the other), provides positional information about the held object. Several studies on multisensory grasping showed that the advantages in multisensory (visuo-haptic) action performance stem from the use of haptic positional information from the contralateral hand (Camponogara, 2023; Camponogara & Volcic, 2019a, 2019b, 2021b, 2022) or a handheld tool (Camponogara, Farnè, & Volcic, 2024) to guide the grasping of objects. Concurrently, in multisensory reaching, haptic positional information supports action performance when vision is suddenly withdrawn (Camponogara & Volcic, 2021a). Thus, a decay in action performance may have been prevented by the use of similar haptic positional information from the arm holding the target and the arm reaching it. This suggests that the decay in the Delay block was due to the lack of direct comparisons between arm postures. Moreover, our results also suggest that when haptic on-line information is not available, actions rely on arm postural information stored in haptic memory.

According to our accuracy results, haptic memory can last up to six seconds, but it is contingent on the arms’ postures. When the postures of the sensing and reaching arms are similar, information can be stored longer compared to when they are not. Reaching previously sensed targets on the left side of the body, indeed, requires the transformation of one posture (sensing arm) to another (reaching arm), which may have affected action performance. It is not to exclude, however, that our results may be related to biomechanical factors concerning the control of different degrees of freedom of the joints when reaching targets in the contralateral (Left) or ipsilateral (Center) hemifields. Reaching targets on the Left may have required a heavier use of the shoulder joint than Center targets, where actions have mainly involved the elbow joint. Thus, reaching contralateral targets may have required a higher level of complexity in hand control than ipsilateral targets. This may have affected action accuracy, especially for targets in the near space (van Beers, Baraduc, & Wolpert, 2002). Further investigations might rule out this alternative explanation by focusing on the role of biomechanical factors in memory-guided haptic reaching.

The importance of our results can be extended beyond the haptic domain, as they may shed some light on the underlying sensorimotor transformation processes in multisensory action performance. According to Tagliabue and McIntyre (2014) and Bernard-Espina, Beraneck, Maier, and Tagliabue (2021), multisensory action guidance could be explained by a set of different models. A concurrent model, where haptic and visual information about the target is processed in parallel with haptic and visual information from the hand performing the action; a fully concurrent model, where haptic and visual information from the target and the hand are also cross-transformed and compared to the sensory estimate of the other sense (i.e., compared with haptics for vision and with vision for haptics); a convergent model, where haptic and visual target information converges with the reaching hand information into a unique multimodal estimate; a fully convergent model, where haptic and visual information from both the target and the hand are independently estimated and compared within a unique multimodal estimate; and, a hybrid convergent/concurrent model where each sensory information from the target and the hand is cross transformed and compared with visual and haptic estimates. Based on our findings and previous work on visually guided reaching (Berkinblit et al., 1995; Darling & Miller, 1993; Heath & Binsted, 2007; Heath et al., 2004; Soechting & Flanders, 1989a, 1989b; Tillery et al., 1991; Westwood et al., 2001, 2003), it is possible that only visual inputs require cross-modal transformation, whereas in the haptic domain, sensing and reaching occur within the same sensory modality.

In conclusion, we demonstrated that memory-guided reaching in the haptic domain is guided by the encoding of target and reaching arm positions through direct comparison of arm postural information. The effect of a delay between the sensing and the reaching phases depends on the postures of the two arms. When arm postures are similar, there is no dramatic decrease in action performance with increasing time delays. Instead, when arm postures differ, action performance degrades as the delay between sensing and reaching increases. It is worth considering that this study is limited to a six-second delay, and further studies could explore longer delays to investigate at what delay actions performed with comparable arm postures are most impaired.

## CRediT author statement

**Ivan Camponogara:** Conceptualization, Methodology, Software, Validation, Formal analysis, Investigation, Data Curation, Writing - Original Draft, Writing - Review & Editing, Visualization. **Robert Volcic:** Conceptualization, Methodology, Software, Validation, Formal analysis, Data Curation, Resources, Writing - Review & Editing, Visualization, Supervision.

## Data availability

All data are available at the following link: https://osf.io/8nb6u.

## Acknowledgements

The authors would like to thank Faisal Abdulhadi for the help in data collection. We acknowledge the support of the NYU Abu Dhabi Research Enhancement Fund (grant RE183), the NYUAD Center for Artificial Intelligence and Robotics, funded by Tamkeen under the NYUAD Research Institute Award CG010, the NYUAD Center for Brain and Health, funded by Tamkeen under the NYUAD Research Institute Award CG012, and, the support of the Zayed University Research Incentive Fund (grant 23076).

## Additional information

### Competing interests

The authors declare no competing interests. Correspondence and requests for materials should be addressed to I.C.

## Notes

https://osf.io/8nb6u

### Competing Interest Statement

The authors have declared no competing interest.

